# Induction of cell death in U2OS human cancer cells by Ta_3_N_5_

**DOI:** 10.1101/2024.10.29.620774

**Authors:** Tomoatsu Nozaki, Kazuto Watanabe, Tomoaki Watanabe, Takahiro J. Nakamura

**Author notes:** Corresponding author, Takahiro J. Nakamura, Ph.D., Laboratory of Animal Physiology, School of Agriculture, Meiji University, 1-1-1 Higashimita, Tama-ku, Kawasaki, Kanagawa 214-8571, Japan.

## Abstract

Photocatalysts are widely used in modern society, with their applications extending into the field of medicine. Ultraviolet irradiation is used for photocatalysis to generate reactive oxygen species (ROS). Ta_3_N_5_ has attracted attention as a next-generation photocatalyst because it produces ROS upon exposure to visible light. Although ROS generation using visible light irradiation is useful in the field of life sciences, the toxicity of Ta_3_N_5_ remains unknown. In the present study, we evaluated the cytotoxicity of Ta_3_N_5_ using U2OS cells in terms of the intensity, irradiation time, and wavelength of light and Ta_3_N_5_ dose. We also investigated the mode and mechanism of cell death induced by Ta_3_N_5_. Cytotoxicity increased with increasing light intensity, irradiation time, and Ta_3_N_5_ dose. In contrast, high-intensity blue light irradiation induced cytotoxicity with or without Ta_3_N_5_, indicating that irradiation to green and red lights, which can selectively toxify Ta_3_N_5_-treated cells, is more useful than irradiation to blue light. Moreover, Ta_3_N_5_ induced secondary necrotic death of U2OS cells. These results suggest that Ta_3_N_5_ can be used as a photocatalyst in medical and life science fields.

## Introduction

A photocatalyst produces photoinduced holes with oxidizing potential and photoinduced electrons with reducing potential, by absorbing light energy from a source such as sunlight. The generated holes react with oxygen and water molecules adsorbed on the surface of the photocatalyst to produce reactive oxygen species (ROS) (Chuah et al., 2017). TiO_2_ is the primarily used photocatalyst. It is widely used for purifying air and water, deodorization, sterilization, and antibacterial effects, and is applied to the exterior and interior walls of buildings (Nozawa et al., 2001; Kühn et al., 2003; Yu et al., 2003; Zhao and Yang, 2003; Vincent et al., 2007; Wu et al., 2009; Ahmed et al., 2011; Di Paola et al., 2012; Zhang et al., 2017). However, its use is associated with toxicological concerns (Vamanu et al., 2008). TiO_2_ has been extensively studied for ecotoxicity, phytotoxicity, cytotoxicity, and genotoxicity (Jośko and Oleszczuk, 2014; Sha et al., 2015; Fytianos et al., 2020). In addition, conventional photocatalysts, including TiO_2_, exhibit oxidative potential when irradiated with ultraviolet (UV) light. UV irradiation is cytotoxic to corneal fibroblasts and inhibits cell proliferation (Chang et al., 2008). Therefore, application of TiO_2_ to living bodies is associated with concerns regarding nonspecific damage upon UV exposure (Onuma et al., 2009).

Ta_3_N_5_ is considered a promising next-generation photocatalytic material since 2000 (Hitoki et al., 2002). Single-crystal Ta_3_N_5_ has been developed in 2018 (Pihosh et al., 2023). Ta_3_N_5_ is characterized by its photocatalytic activity upon absorption of light within the blue–green range (400–600 nm, visible light). Therefore, Ta_3_N_5_ does not require UV irradiation for the photocatalytic activity, and its use may be relatively less damaging to living organisms (Hitoki et al., 2002). Extensive research has been conducted on the material productions, applications, and properties of Ta_3_N_5_ (e.g., Pihosh et al., 2020; Xiao et al., 2020; Xiao et al. 2022), however, its cytotoxicity and the form of cell death induced by Ta_3_N_5_ remain largely unexplored, thereby hindering its biological applications.

In the present study, we evaluated the cytotoxicity of Ta_3_N_5_ using a human osteosarcoma cell line, U2OS, in terms of the intensity, irradiation time, and wavelength of light and the Ta_3_N_5_ dose, and investigated the mode and mechanism of cell death induction. The tumor suppressor gene p53 responds to DNA damage or cellular stress as a transcription factor and trans-activates or represses several target genes that induce cell cycle arrest or apoptosis. We selected U2OS cells for this study because they possess wild-type p53 and have been used in various biomedical studies, such as analyses of apoptosis and DNA repair pathways (Greenblatt et al., 1994; Samuels-Lev et al., 2001).

## Materials and Methods

### Light irradiation

Six light-emitting diode panels capable of emitting blue (460–470 nm), green (520–525 nm), and red (620–625 nm) lights were mounted on an aluminum plate, and light intensity was adjusted after wiring. A light-irradiation device was fabricated with a stand at a certain distance from the object for its even irradiation. The light intensity was set to 30, 60, and 90 mW/cm^2^. Light intensities were measured using a radiation meter LP1 (Sanwa Denki Keiki, Tokyo, Japan). The irradiation device was placed inside a CO_2_ incubator (Astec, Fukuoka, Japan) and used for the experiments.

### Cell culture

U2OS cells were purchased from ATCC (VA, USA). The cells are grown in Dulbecco’s Modified Eagle Medium (DMEM) (Thermo Fisher Scientific, MA, USA) supplemented with 10% fetal bovine serum (FBS) (Nichirei, Tokyo, Japan) and 100 U/mL penicillin–streptomycin (Thermo Fisher Scientific). For deactivation, cells were cultured in DMEM containing 4.5 g/L D-glucose, L-glutamine (Thermo Fisher Scientific) and 10% FBS.

### Preparation of Ta_3_N_5_ suspension

Ta_3_N_5_ powder (0.5 g; Kozyundo Chemical Laboratories, Saitama, Japan) was added to 5 mL phosphate-buffered saline (PBS) to prepare a 0.1 g/mL Ta_3_N_5_ suspension, and the suspension was autoclaved.

### Trypan blue staining

U2OS cells were seeded at 1.5 × 10^4^ cells/well in 8-well chamber slides (Matsunami Glass Industry, Osaka, Japan) and incubated for 1 day. Thereafter, 0, 1.6, or 3.2 mg/mL Ta_3_N_5_ was added to the green light-irradiated cell groups at a ratio of culture medium to Trypan blue solution (Thermo Fisher Scientific) of 1:1, and cells were stained, washed, and sealed. Images of cells were captured using an IX-53 camera-equipped biological microscope (Olympus, Tokyo, Japan).

### Measurement of viable cells in the Ta_3_N_5_-treated group

U2OS cells were seeded at 3 × 10^4^ cells/well in 48-well plates (Violamo, Osaka, Japan). After 1 day of culture, Ta_3_N_5_ was added at 0, 1.6, or 3.2 mg/mL concentrations to 16 wells each, and cells were irradiated with blue, green, and red lights for 1, 2, and 4 h. Cell viability was measured using a WST-8 assay. Specifically, after washing cells with 100 μL PBS, 100 μL DMEM (without phenol red) and 10 μL Cell Counting Kit-8 (Dojindo Laboratories, Kumamoto, Japan) were added to each well, and the plates were incubated at 37 °C for 3 h. Then, 80 μL supernatant was transferred to a 96-well plate (Violamo), and the absorbance at 450 nm was measured using an Enspire 2300 spectrophotometer (Perkin Elmer, MA, USA) to determine the number of viable cells. The absorbance values were converted to a percentage, with the absorbance of the non-illuminated group considered as 100, and relative cell viability was calculated.

### Identification of cell death pathways induced by Ta_3_N_5_ using fluorescence immunostaining

Fluorescence immunostaining was performed to analyze the morphology of apoptotic cells. U2OS cells were seeded at 1.5 × 10^4^ cells/well in 8-well chamber slides (Matsunami Glass Industry) and cultured for 1 day. Then, 20 μL of 3.2 mg/mL Ta_3_N_5_ was added to each well, and the slides were irradiated with 90 mW/cm^2^ green light for 1 h. After irradiation, cells were washed with PBS, stained using a MEBCYTO Apoptosis Kit (Annexin V-FITC Kit; MBL, Nagoya, Japan), according to the manufacturer’s protocol. Fluorescence was observed using a confocal laser scanning microscope (LSM 880; Zeiss, Oberkochen, Germany).

Images were processed using ImageJ (National Institutes of Health, Bethesda, MD, USA) to quantify the size of nuclei. For the group not treated with Ta_3_N_5_, cells were stained with 4’,6-diamidino-2-phenylindole (DAPI), and images were captured using a confocal laser scanning microscope.

### Detection of DNA fragmentation in dead cells

U2OS cells (1.0 × 10^7^) were seeded in a 6-mm dish. After 24 h, cells were harvested using a cell scraper. DNA was isolated, and a DNA ladder assay was performed using an ApopLadder Ex kit (Takara Bio, Shiga, Japan), according to the manufacturer’s instructions. Extracted DNA was dissolved in 10 μL of 1× Tris-EDTA buffer, mixed with 2 μL of 6× loading buffer and 1 μL of Midori Green Direct (Nippon Genetics, Tokyo), and electrophoresed on a 2% agarose gel for visualization. Images were acquired using a Limited-STAGE II gel imaging system (Ams System Science, Osaka, Japan).

### Statistics analysis

Results of cytotoxicity assay and nuclear size are expressed as mean ± standard error. One-way analysis of variance (ANOVA) was used to compare three or more groups. A *P*-value < 0.05 was considered statistically significant. In case of a significant difference, multiple comparison with one-way ANOVA followed by Tukey’s test was used to test the significance of the data. Outliers were excluded using the Smirnoff–Grubbs test.

## Results

### Cytotoxicity induced by Ta_3_N_5_

Most cells in the control group (without Ta_3_N_5_ treatment) were live as they were not stained with trypan blue. However, Ta_3_N_5_ doses of 1.6 and 3.2 mg/mL and irradiation with 90 mW/cm^2^ green light for 4 h resulted in considerable cell death (Figure 1). A decrease in cell density was observed in the treated group owing to cell death and detachment of dead cells during washing. Similarly, the shape of dead cells was changed, with healthy cells being polygonal and dead cells being spherical.

**Fig. 1.**
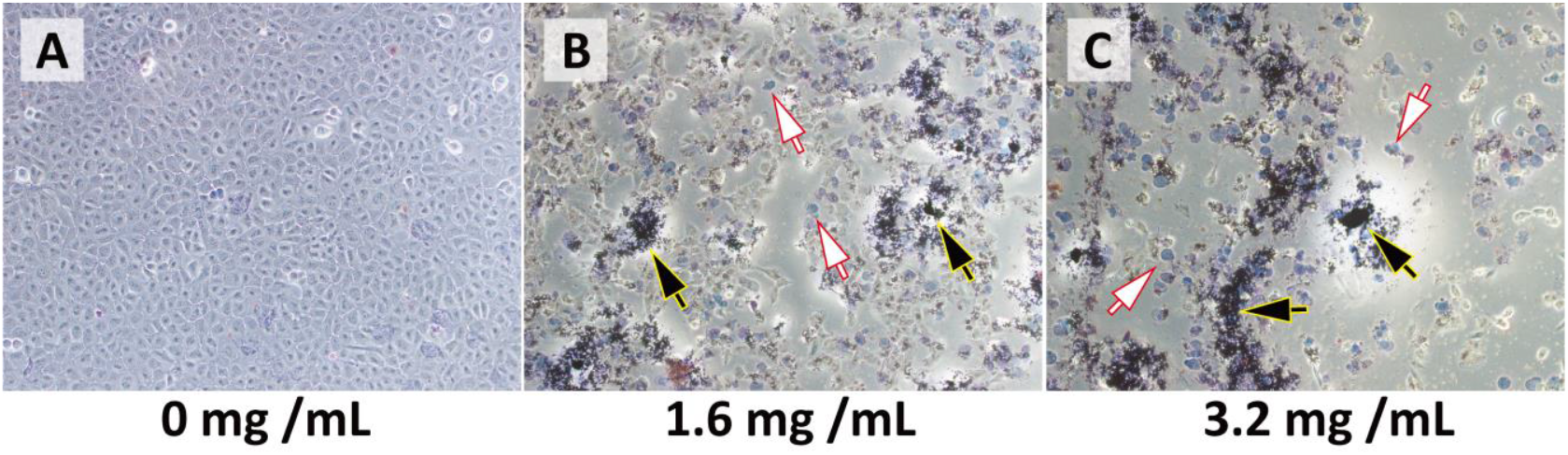
Images of trypan blue-stained cells showing Ta_3_N_5_-induced cytotoxicity. Trypan blue staining during irradiation with green light at 90 mW/cm^2^ for 4 h. (A) Cells without Ta_3_N_5_ treatment. (B) Cells treated with 1.6 mg/mL Ta_3_N_5_. (C) Cells treated with 3.2 mg/mL Ta_3_N_5_. Blue-stained cells (white arrows) and aggregated Ta_3_N_5_-treated cells (black arrows).

### Characterization of cytotoxicity induced by Ta_3_N_5_

The cytotoxicity of 3.2 mg/mL Ta_3_N_5_ gradually increased with increasing light intensities (at 30, 60, and 90 mW/cm^2^) at a fixed irradiation time of 4 h; viable cells were 23.1 ± 0.5%, 20.2 ± 0.1%, and 3.6 ± 0.0% for blue light irradiation, 58.6 ± 0.7%, 42.3 ± 0.5%, and 35.4 ± 1.2% for green light irradiation, and 64.7 ± 0.5%, 61.6 ± 1.1%, and 30.6 ± 0.6% for red light irradiation, respectively (*P* < 0.01; Figure 2A).

**Fig. 2.**
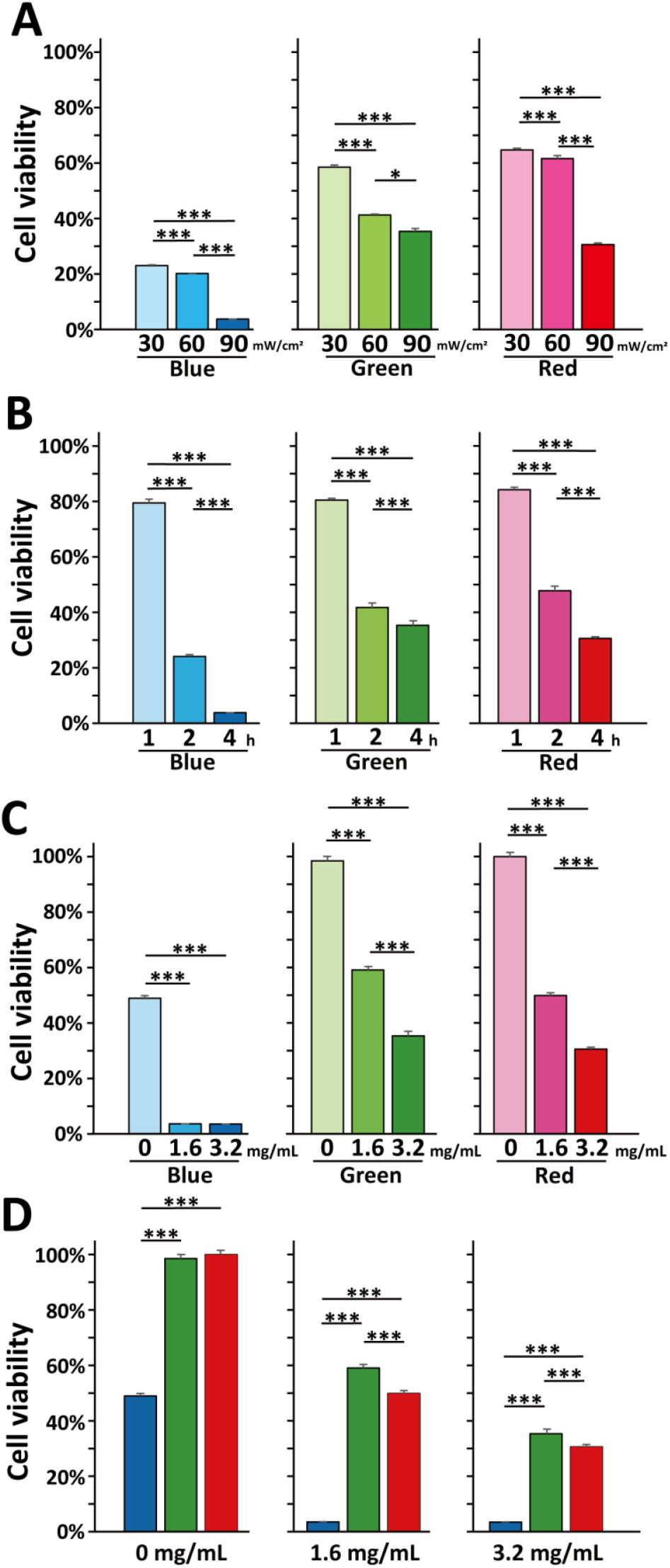
Light intensity, irradiation time, Ta_3_N_5_ dose, and wavelength dependence of Ta_3_N_5_ cytotoxicity on U2OS cells. (A) Viability of cells treated with 3.2 mg/mL Ta_3_N_5_ and exposed to blue, green, or red lights at intensities of 30, 60, or 90 mW/cm^2^ for 4 h. (B) Viability of cells treated with 3.2 mg/mL Ta_3_N_5_ and exposed to 90 mW/cm^2^ blue, green, or red lights for 1, 2, or 4 h. (C) Viability of cells treated with 0, 1,6, or 3.2 mg/mL Ta_3_N_5_ and exposed to 90 mW/cm^2^ blue, green, or red lights for 4 h. (D) Viability of cell treated with 0, 1,6, or mg/mL Ta_3_N_5_ and exposed to 90 mW/cm^2^ of blue, green, or red lights for 4 h. \**P* < 0.05, ** *P* < 0.01, *** *P* < 0.001, Tukey’s test.

Similarly, the cytotoxicity of 3.2 mg/mL Ta_3_N_5_ at 90 mW/cm^2^ light intensity gradually increased with irradiation time (1, 2, and 4 h); viable cells were 77.5 ± 1.4%, 24.1 ± 0.7%, and 3.6 ± 0.0% for blue light irradiation, 80.4 ± 0.9%, 41.8 ± 1.2%, and 35.4 ± 1.2% for green light irradiation, and 84.3 ± 0.9%, 47.8 ± 1.7%, and 30.6 ± 0.6% for red light irradiation, respectively (*P* < 0.01; Figure 2B).

At Ta_3_N_5_ doses of 0, 1.6, and 3.2 mg/mL and a light intensity of 90 mW/cm^2^ for 4 h, viable cells were 49.0 ± 0.9%, 3.6 ± 0.0%, and 3.6 ± 0.0% for blue light irradiation, 98.5 ± 1.2%, 59.1 ± 0.9%, and 35.4 ± 1.2% for green light irradiation, and 100.0 ± 1.1%, 49.9 ± 1.0%, and 30.6 ± 0.6% for red light irradiation, respectively (*P* < 0.01; Figure 2C), indicating a dose-dependent increase in cytotoxicity of Ta_3_N_5_.

Blue light exhibited the highest cytotoxicity at 3.2 mg/mL Ta_3_N_5_ dose at an intensity of 90 mW/cm^2^ for 4 h (*P* < 0.01; Figure 2D). In addition, blue light of relatively high intensity considerably killed cells without Ta_3_N_5_ treatment.

### Evaluation of cell death pathways using fluorescence immunostaining

Fluorescence immunostaining with propidium iodide (PI) and Annexin V showed that majority of cells in the control group were not stained with PI and Annexin V, indicating no cellular apoptosis (Figure 3A). Treatment with staurosporine, which induces apoptosis, caused severe cell death as indicated by cells stained with Annexin V or both Annexin V and PI. Similarly, Ta_3_N_5_-treated cells were stained with Annexin V or both Annexin V and PI, thereby indicating cellular apoptosis.

**Fig. 3.**
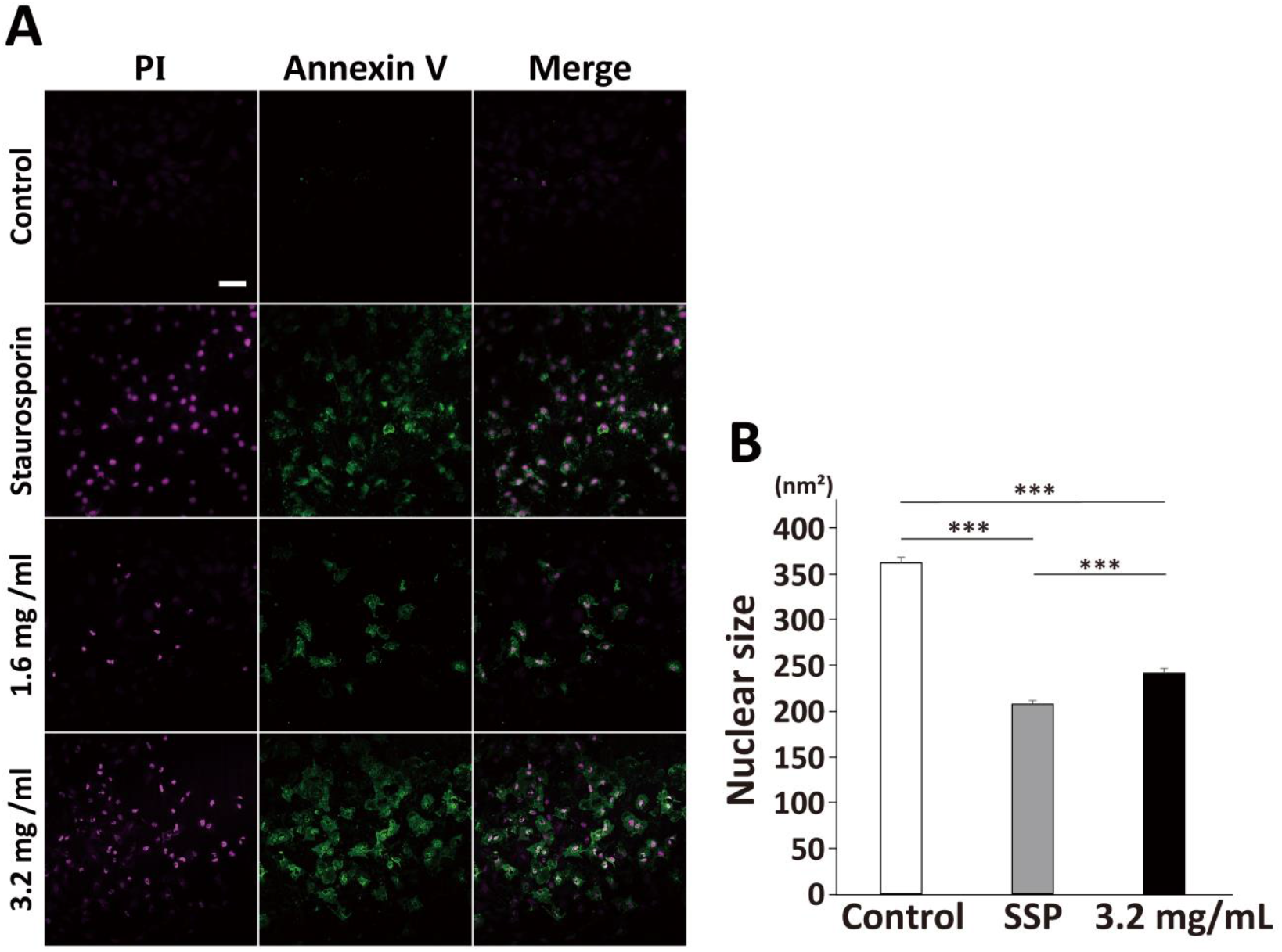
PI and Annexin V staining and size of nuclei of cells subjected to Ta_3_N_5_-induced cytotoxicity. Annexin V and PI staining of cells irradiated with 90 mW/cm^2^ green light for 1 h. Green fluorescence indicates the exposure of phosphatidylserine outside the plasma membrane, and red fluorescence indicates the nucleus. From left to right, images of PI- and Annexin V-stained cells, and superimposed images, for the control, staurosporine (SSP)-treated, 1.6 mg/mL Ta_3_N_5_-treated, and 3.2 mg/mL Ta_3_N_5_-treated groups, from top to bottom. Scale bar, 100 nm. (B) Image J analysis of nuclear size. \*\*\**P* < 0.001, Tukey’s test.

The nuclei of cells undergoing apoptosis were significantly reduced in size (control group, 362.1 ± 6.3 nm^2^; staurosporine-treated group, 207.3 ± 4.1 nm^2^; 3.2 mg/mL Ta_3_N_5_-treated group group, 241.8 ± 4.9 nm^2^) (*P* < 0.01; Figure 3B).

### Evaluation of cell death pathways

We next evaluated DNA fragmentation, a characteristic aspect of apoptosis resulting from the activation of endogenous endonuclease (Figure 4). DNA of staurosporine-treated cells showed clean ladder-like bands, confirming that the method was suitable for characterizing apoptosis. DNA fragmentation was not observed in the control group, and only one polymeric band was observed. In contrast, in the Ta_3_N_5_-treated group, smeared bands were noticed for all periods of light irradiation, and no significant differences among them were observed.

**Fig. 4.**
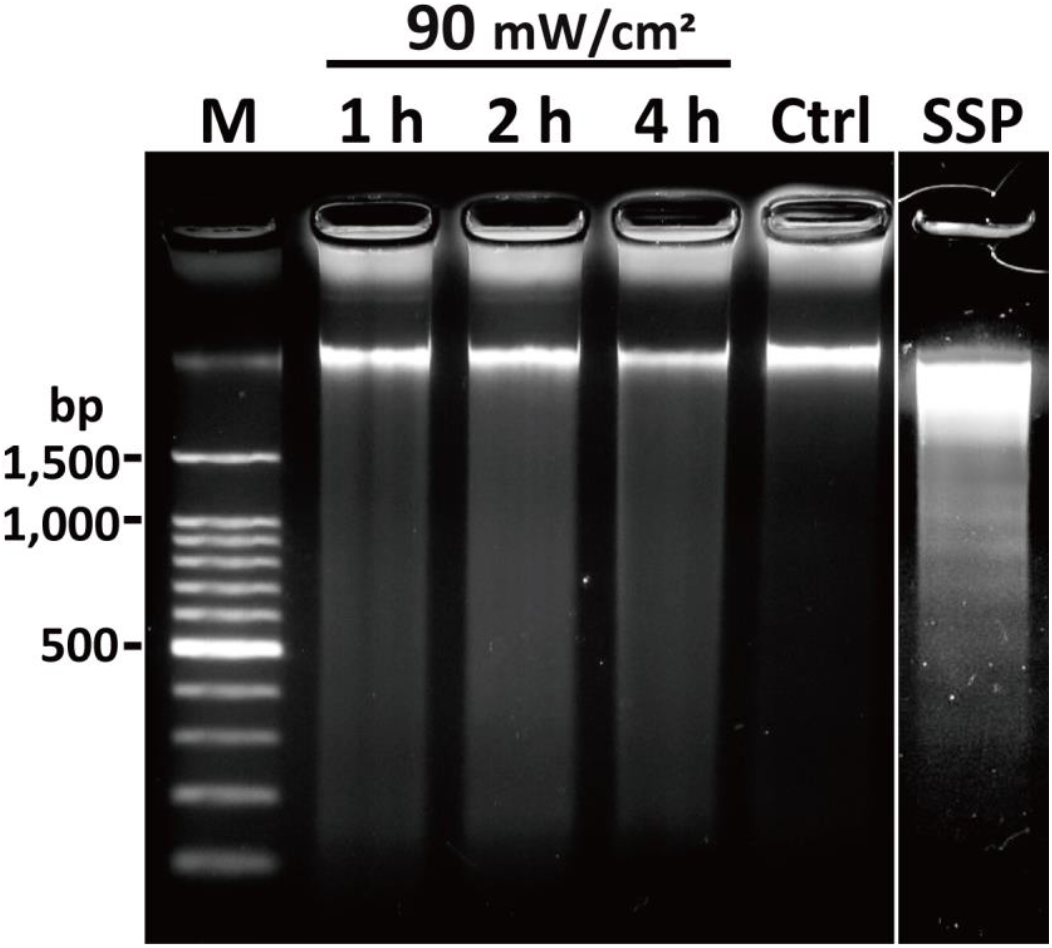
Gel image of fragmented DNA after Ta_3_N_5_ treatment. Representative image of fragmented DNA of Ta_3_N_5_-treated cells. The image on the right shows a ladder of fragmented DNA extracted from staurosporine-treated cells. From left to right, DNA extracted from cells irradiated using 90 mW/cm^2^ green light for 1, 2, or 4 h, and that of the control group. Ctrl, control; SSP, staurosporine.

## Discussion

In the present study, we found that Ta_3_N_5_ exhibited irradiation time- and dose-dependent cytotoxicity and induced secondary necrotic death of U2OS cells. Therefore, Ta_3_N_5_ can be a useful photocatalyst in the fields of medical and life sciences. TiO2 is a metal nitride with photocatalytic properties, which has been widely used in the chemical field, and its utilization has been extended further to the medical fields (Jafari et al., 2020). Several studies on the toxicity of TiO2 have been conducted (Jośko and Oleszczuk, 2014; Sha et al., 2015; Fytianos et al., 2020). Ta_3_N_5_ is a recently developed photocatalyst, which shows photocatalytic effect in the visible wavelengths of light (Pihosh et al., 2023). It has attracted attention as a substitute for TiO2 that requires UV irradiation. However, Ta_3_N_5_ has not been compared with TiO2 for use in the medical and physiological fields.

Similar to that by TiO_2_ (Hamzeh and Sunahara, 2013), Ta_3_N_5_ exhibited irradiation time- and dose-dependent cytotoxicity. Unlike that by green and red lights, blue light irradiation at high intensity induced cell death even in the absence of Ta_3_N_5_, probably owing to the high energy of blue light that has a wavelength close to that of UV light. Blue light induces apoptosis via excessive production of intracellular ROS and a decrease in mitochondrial membrane potential (Hao et al., 2023), and the same phenomenon may have occurred in the group not treated with Ta_3_N_5_. This suggests that irradiation using green and red lights can induce cytotoxicity specifically in the Ta_3_N_5_-treated group. Cytotoxicity under green- and red-light irradiation is unique to Ta_3_N_5_ and has not been reported in toxicity studies using other photocatalysts. Therefore, if Ta_3_N_5_ could be accumulated in target cells, such as cancer cells, or in a tissue-specific manner, it can induce toxicity in specific cells without harming normal cells.

TiO_2_ induces apoptosis and necrosis in human monoblast U937 and mouse lymphocytic leukocyte L1210 cells (Vamanu et al., 2008; Takaki et al., 2014). In the present study, we investigated whether Ta_3_N_5_ induces apoptosis and/or necrosis. Apoptotic cells expose phosphatidylserine to the plasma membrane exterior as an “eat me” signal (Li, 2012). Therefore, apoptotic cells can be visualized using fluorescein isothiocyanate-modified Annexin V, which specifically binds to phosphatidylserine and emits green fluorescence. In the Ta_3_N_5_-treated group, several cells emitted green fluorescence owing to Annexin V binding. The measurement of nuclear size indicated nuclear shrinkage, which is one of a hallmark of apoptosis. Cleavage of DNA, approximately every 200 bp, during the early stages of apoptosis may prevent the transfer of potentially active genetic material to neighboring cells, as the apoptotic bodies are phagocytosed by macrophages (Arends et al., 1990). In contrast, necrosis is characterized by nonspecific DNA cleavage and observation of smeared bands on gel (Schumer et al., 1992). In our study, DNA ladders were observed for staurosporine-treated cells, whereas smear-like bands were noticed for Ta_3_N_5_-treated cells. Therefore, Ta_3_N_5_ may have induced secondary necrosis in U2OS cells.

Ta_3_N_5_ produces ROS upon light irradiation. ROS might have induced oxidative stress-responsive apoptosis via the mitogen-activated protein kinase pathway or Ta_3_N_5_ might have induced apoptosis by lowering the mitochondrial membrane potential or causing DNA damage (Nakano and Vousden, 2001; Oda et al., 2000; Green, 2010; Lee et al., 2014). However, in our *in vitro* experimental system, the absence of macrophages to phagocytose apoptotic cells may have caused the final rupture of the membrane, resulting in a morphology similar to that of necrotic cells. The mechanism of this type of cell death is still unclear. Moreover, the results for cell death induced by nanomaterials are often inconsistent and even contradictory because of the differences in the shape and surface chemistry of nanomaterials, experimental design and conditions, and several other factors. The specific molecular mechanisms of Ta_3_N_5_-induced cell death, including the expression of caspases and apoptosis-related genes, need to be investigated. In addition, *in vivo* cytotoxicity and cell death patterns associated with Ta_3_N_5_ need to be explored.

TiO_2_ is a metal nitride with photocatalytic properties, which has been widely used in the chemical field, and its utilization has been extended further to the medical fields (Jafari et al., 2020). Several studies on the toxicity of TiO_2_ have been conducted (Jośko and Oleszczuk, 2014; Sha et al., 2015; Fytianos et al., 2020). Ta_3_N_5_ is a recently developed photocatalyst, which shows photocatalytic effect in the visible wavelengths of light (Pihosh et al., 2023). It has attracted attention as a substitute for TiO_2_ that requires UV irradiation. However, Ta_3_N_5_ has not been compared with TiO_2_ for use in the medical and physiological fields.

## Acknowledgments

This work was supported by Japan Society for the Promotion of Science (JSPS) KAKENHI (grant number 19K06360 to T.J.N.) and Iketani Science and Technology Foundation (grant number 0341219-A to T.J.N.). We thank Editage (www.editage.jp) for English language editing.

## References

Ahmed, S., Rasul, M.G., Brown, R. and Hashib, M.A. (2011): Influence of parameters on the heterogeneous photocatalytic degradation of pesticides and phenolic contaminants in wastewater: A short review. J. Environ. Manage., 92, 311–330.

Arends, M.J., Morris, R.G. and Wyllie, A.H. (1990): Apoptosis: The role of the endonuclease. Am. J. Pathol., 136, 593–608.

Chang, S.-W., Chou, S.-F. and Chuang, J.-L. (2008): Mitomycin C potentiates ultraviolet-related cytotoxicity in corneal fibroblasts. Cornea., 27, 686–692.

Chuah, X.-F., Lee, K.-T., Cheng, Y.-C., Lee, P.-F. and Lu, S.-Y. (2017): Ag/AgFeO2: An outstanding magnetically responsive photocatalyst for HeLa cell eradication. ACS Omega., 2, 4261–4268.

Di Paola, A., García-López, E., Marcì, G. and Palmisano, L. (2012): A survey of photocatalytic materials for environmental remediation. J. Hazard. Mater., 211-212, 3–29.

Fytianos, G., Rahdar, A. and Kyzas, G.Z. (2020): Nanomaterials in cosmetics: Recent updates. Nanomaterials (Basel)., 10, 979.

Green, D.R. (2010): Means to an End: Apoptosis and Other Cell Death Mechanisms. Cold Spring Harbor Laboratory Press, USA.

Greenblatt, M.S., Bennett, W.P., Hollstein, M. and Harris, C.C. (1994): Mutations in the p53 tumor suppressor gene: clues to cancer etiology and molecular pathogenesis. Cancer Res., 54, 4855–4878.

Hamzeh, M. and Sunahara, G.I. (2013): In vitro cytotoxicity and genotoxicity studies of titanium dioxide (TiO_2_) nanoparticles in Chinese hamster lung fibroblast cells. Toxicol. In Vitro., 27:864–873.

Hao, W., Zhao, C., Li, G., Wang, H., Li, T., Yan, P. and Wei, S. (2023): Blue LED light induces cytotoxicity via ROS production and mitochondrial damage in bovine subcutaneous preadipocytes. Environ. Pollut., 322, 121195.

Hitoki, G., Ishikawa, A., Takata, T., Kondo, J.N., Hara, M. and Domen, K. (2002): Ta3N5 as a novel visible light-driven photocatalyst (λ < 600 nm). Chem. Lett., 31, 736–737.

Jafari, S., Mahyad, B., Hashemzadeh, H., Janfaza, S., Gholikhani, T. and Tayebi, L. (2020): Biomedical applications of TiO2 nanostructures: Recent advances. Int. J. Nanomedicine., 15, 3447–3470.

Jośko, I. and Oleszczuk, P. (2014): Phytotoxicity of nanoparticles--problems with bioassay choosing and sample preparation. Environ. Sci. Pollut. Res. Int., 21, 10215–10224.

Kühn, K.P., Chaberny, I.F., Massholder, K., Stickler, M., Benz, V.W., Sonntag, H.-G. and Erdinger, L. (2003): Disinfection of surfaces by photocatalytic oxidation with titanium dioxide and UVA light. Chemosphere., 53, 71–77.

Lee, H.-S., Hwang, C.Y., Shin, S.-Y., Kwon, K.-S. and Cho, K.-H. (2014): MLK3 is part of a feedback mechanism that regulates different cellular responses to reactive oxygen species. Sci. Signal., 7, ra52.

Li, W. (2012): Eat-me signals: Keys to molecular phagocyte biology and “appetite” control. J. Cell. Physiol., 227, 1291–1297.

Nakano, K. and Vousden, K.H. (2001): PUMA, a novel proapoptotic gene, is induced by p53. Mol. Cell. 7, 683–694.

Nozawa, M., Tanigawa, K., Hosomi, M., Chikusa, T. and Kawada, E. (2001): Removal and decomposition of malodorants by using titanium dioxide photocatalyst supported on fiber activated carbon. Water Sci. Technol., 44, 127–133.

Oda, E., Ohki, R., Murasawa, H., Nemoto, J., Shibue, T., Yamashita, T., Tokino, T., Taniguchi, T. and Tanaka, N. (2000): Noxa, a BH3-only member of the Bcl-2 family and candidate mediator of p53-induced apoptosis. Science., 288, 1053–1058.

Onuma, K., Sato, Y., Ogawara, S., Shirasawa, N., Kobayashi, M., Yoshitake, J., Yoshimura, T., Iigo, M., Fujii, J. and Okada, F. (2009): Nano-scaled particles of titanium dioxide convert benign mouse fibrosarcoma cells into aggressive tumor cells. Am. J. Pathol., 175, 2171–2183.

Pihosh, Y., Nandal, V., Higashi, T., Shoji, R., Bekarevich, R., Nishiyama, H., Yamada, T., Nicolosi, V., Hisatomi, T., Matsuzaki, H., Seki, K. and Domen, K. (2023): Tantalum nitride-enabled solar water splitting with efficiency above 10%. Adv. Energy Mater., 13, 2301327.

Pihosh, Y., Minegishi, T., Nandal, V., Higashi, T., Katayama, M., Yamada, T., Sasaki, Y., Seki, K., Suzuki, Y., Nakabayashi, M., Sugiyama, M., Domen, K. (2020): Ta3N5-Nanorods enabling highly efficient water oxidation via advantageous light harvesting and charge collection. Energy Environ. Sci., 13, 1519.

Samuels-Lev, Y., O’Connor, D.J., Bergamaschi, D., Trigiante, G., Hsieh, J.K., Zhong, S., Campargue, I., Naumovski, L., Crook, T. and Lu, X. (2001): ASPP proteins specifically stimulate the apoptotic function of p53. Mol. Cell., 8, 781–794.

Schumer, M., Colombel, M.C., Sawczuk, I.S., Gobé, G., Connor, J., O’Toole, K.M., Olsson, C.A., Wise, G.J. and Buttyan, R. (1992): Morphologic, biochemical, and molecular evidence of apoptosis during the reperfusion phase after brief periods of renal ischemia. Am. J. Pathol., 140, 831–838.

Sha, B., Gao, W., Cui, X., Wang, L. and Xu, F. (2015): The potential health challenges of TiO2 nanomaterials. J. Appl. Toxicol. 35, 1086–1101.

Takaki, K., Higuchi, Y., Hashii, M., Ogino, C. and Shimizu, N. (2014): Induction of apoptosis associated with chromosomal DNA fragmentation and caspase-3 activation in leukemia L1210 cells by TiO2 nanoparticles. J. Biosci. Bioeng., 117, 129–133.

Vamanu, C.I., Cimpan, M.R., Høl, P.J., Sørnes, S., Lie, S.A. and Gjerdet, N.R. (2008) Induction of cell death by TiO2 nanoparticles: studies on a human monoblastoid cell line. Toxicol. In Vitro., 22, 1689–1696.

Vincent, G., Queffeulou, A., Marquaire, P.M. and Zahraa, O. (2007): Remediation of olfactory pollution by photocatalytic degradation process: Study of methyl ethyl ketone (MEK). J. Photochem. Photobiol. A Chem., 191, 42–50.

Wu, R.-J., Chen, C.-C., Chen, M.-H. and Lu, C.-S. (2009): Titanium dioxide mediated heterogeneous photocatalytic degradation of gaseous dimethyl sulfide: Parameter study and reaction pathways. J. Hazard. Mater., 162, 945–953.

Xiao, Y., Feng, C., J. Fu, J., F. Wang, F., Li, C., Kunzelmann, V. F., Jiang, C. M., Nakabayashi, M., Shibata, N., Sharp, I. D., Domen, K., Li, Y. (2020): Band structure engineering and defect control of Ta3N5 for efficient photoelectrochemical water oxidation. Nat. Catal., 3, 932.

Xiao, Y., Fan, Z., Nakabayashi, M., Li, Q., Zhou, L., Wang, Q., Li, C., Shibata, N., Domen, K. Li, Y. (2022): Decoupling light absorption and carrier transport via heterogeneous doping in Ta3N5 thin film photoanode. Nat. Commun., 13, 7769.

Yu, J.C., Ho, W., Lin, J., Yip, H. and Wong, P.K. (2003): Photocatalytic activity, antibacterial effect, and photoinduced hydrophilicity of TiO2 films coated on a stainless steel substrate. Environ. Sci. Technol., 37, 2296–2301.

Zhang, X., Xiao, G., Wang, Y., Zhao, Y., Su, H. and Tan, T. (2017): Preparation of chitosan-TiO2 composite film with efficient antimicrobial activities under visible light for food packaging applications. Carbohydr. Polym. 169, 101–107.

Zhao, J. and Yang, X.D. (2003): Photocatalytic oxidation for indoor air purification: a literature review. Build. Environ., 38, 645–654.

